# A Novel Conditional Adra2a-Knockout Mouse Line Reveals Cell-specific Contributions to Specific Dimensions of Sedation

**DOI:** 10.64898/2026.04.14.712766

**Authors:** Noah L. Fryou, Timothy Jiang, Nathan Frick, Paula Kwasniewska, Mikhail Y. Lipin, Max B. Kelz, Steven A. Thomas, Andrew McKinstry-Wu

## Abstract

**Introduction:** Here, we create a conditional *Adra2a* line and use it to show that sedative, hypnotic, and hypothermic effects of α_2_-agonists are neuronally mediated via the α_2A_ adrenergic receptor.

**Methods:** We generated mice with loxP sites flanking *Adra2a* using CRISPR/Cas9 gene targeting. This line was crossed with lines encoding Cre recombinase (Cre) under the control of the Vgat, Snap25, and Dbh promoters. Cell-specific knockout was confirmed using fluorescent in-situ hybridization demonstrating targeted reduction in *Adra2a* mRNA in the appropriate cell types. Mice were given intraperitoneal dexmedetomidine (0.3 or 1 mg/kg) or saline, and 20 minutes later righting reflex was assessed, followed by 3 rounds of rotarod testing, with fall time and end temperature recorded. Spontaneous activity was recorded using beam break for an hour after. Mice of each genotype were implanted with EEG leads and recorded while given 0.3 mg/kg IP dexmedetomidine.

**Results:** We created a conditional knockout and demonstrated cell-type specific reduction of *Adra2a* mRNA in crossed lines with cell-specific Cre. The pan-neuronal *Adra2a* knockout showed resistance to all temperature, sedative, and hypnotic effect endpoints in response to the α_2_-agonist dexmedetomidine. Adrenergic knockout demonstrated resistance to α_2_-agonist hypnosis and moderate resistance to hypothermia and impaired coordination with forced movement. GABAergic knockout showed resistance only to impairment of spontaneous movement by α_2_-agonists. Spectral analysis of the EEG showed an increase in proportion of delta power with a sedative dose of dexmedetomidine in all lines except the pan-neuronal *Adra2a* knockout.

**Discussion:** Future studies will pursue both the specific subtype(s) and location of neuronal populations responsible for sedative, hypnotic, and hypothermic effects of α_2_-agonists.

## Introduction

The α_2A_ adrenergic receptor, encoded in mice by the *Adra2a* gene, plays a crucial role in a variety of critical physiology processes both in the central nervous system (CNS) and throughout the body. Consequently, this receptor is the relevant molecular target for drugs treating conditions in the psychiatric, cardiovascular, ophthalmic, immune, pain, and critical care fields. In the CNS, α_2A_ receptors are the most common α_2_ receptor subtype– found on adrenergic and non-adrenergic neurons as well as astrocytes and microglia.^1^ Medications targeting these central α_2A_ receptors are used to treat ADHD, agitation in individuals with bipolar and schizophrenia, and PTSD, as well as to provide sedation in the ICU and analgesia. Ophthalmically, ciliary epithelium express α_2A_ receptors which are the targets for medications treating glaucoma.^2^ In the periphery, α_2A_ receptors are widespread, found on platelets, adipose tissue, vascular smooth muscle, and macrophages. Accordingly, α_2_ agonist medications are also used in the treatment of hypertension and systemic inflammation. Animal lines with inactivated or knocked-out *Adra2a* have advanced our understanding of adrenergic contributions,^3–5^ specifically the role of signaling through the α_2A_ adrenergic receptor, to these various processes and disease states. However, the distributed expression of this gene, often on multiple different portions of the same system, has limited the use of global knockouts to produce unambiguous answers to key questions for these fields. Certain CNS-mediated effects of α_2_ agonists are a good example of ambiguity that could be resolved with cell-selective knockouts. Hypothermia, hypnosis, and sedation are all known to be central α_2A_-adrenergic-receptor-mediated effects of α_2_ agonists. ^4–8^ The many cell populations in the CNS on which α_2A_ receptors are expressed, however, makes it difficult to discern the relevant site of action. *Adra2a* is present in the CNS on neurons as well as glia, both microglia and astrocytes.^9,10^ Recent evidence has demonstrated central roles for astrocytes in norepinephrine-dependent network activity coordination and synapse remodeling as well as a role in recovery from sedative-hypnotic drug effect,^11–13^ leaving open the possibility that of α_2_ agonists are producing their central effects non-neuronally. On neurons, of α_2A_ receptors are not only present presynaptically on adrenergic neurons, but also on multiple neuronal subtypes postsynaptically.^14,15^ Which of these various populations are responsible for any or all of these effects remains unknown. Here, we described the creation a conditional *Adra2a* mouse line, and the use of that line to produce cell-specific knockouts of the *Adra2a* gene to interrogate the contributions of specific cell populations to centrally-mediated effects of α_2_ agonists.

## Methods

### Mice

All studies were approved by the Institutional Animal Care and Use Committee at the University of Pennsylvania and were conducted in accordance with National Institutes of Health guidelines. Dbh^IRES-Cre^,^16^ Snap25^IRES2-Cre-D^ (RRID:IMSR_JAX:023525),^17^ and Vgat^IRES-Cre^ (RRID:IMSR_JAX:016962)^18^ mouse lines were crossed with the newly-generated floxed Adra2a line (described below) to create cell-line-specific knockouts. All mice were housed on a reverse 12h:12h light/dark cycle (ZT0/lights on at 7:00 PM) with *ad libitum* access to food and water. Adult mice of both sexes between 10 and 45 weeks were used for all assays. Mice had at least 2 weeks of recovery after surgery and 48 hours of rest between experiments.

### *Generation of* Adra2a^fl/fl^ *Mouse Line*

In cooperation with the CRISPR/Cas9 Mouse Targeting Core (University of Pennsylvania) flanking loxP sites were introduced about the single exon of *Adra2a*, the gene encoding the α_2A_ adrenergic receptor, using CRISPR/Cas9 (flanking sgRNA and Cas9mRA microinjected in 1-cell C57/B6J mouse embryos) using established methods.^19,20^ In order to minimize off-target recombination events, we optimized the design of the guide sequences to target within the 5’ UTR and 3’ UTR of the *Adra2a* exon using the CRISPOR program.^21^

Based on that optimization we chose the following 5’ and 3’ targeting sequences:

Adra2a 5’ Target Sequence (20N+PAM; 23N): AGCTAGATCGGTGTACCGCTGGG

Adra2a 3’ Target Sequence (20N+PAM; 23N): AAGGCTACTGGCTTACTTAGGGG

After implantation of transformed embryos, resultant pups were screened for successful insertion of flanking loxP sequences or deletion events via PCR-based genotyping using standard protocols with the following primers:

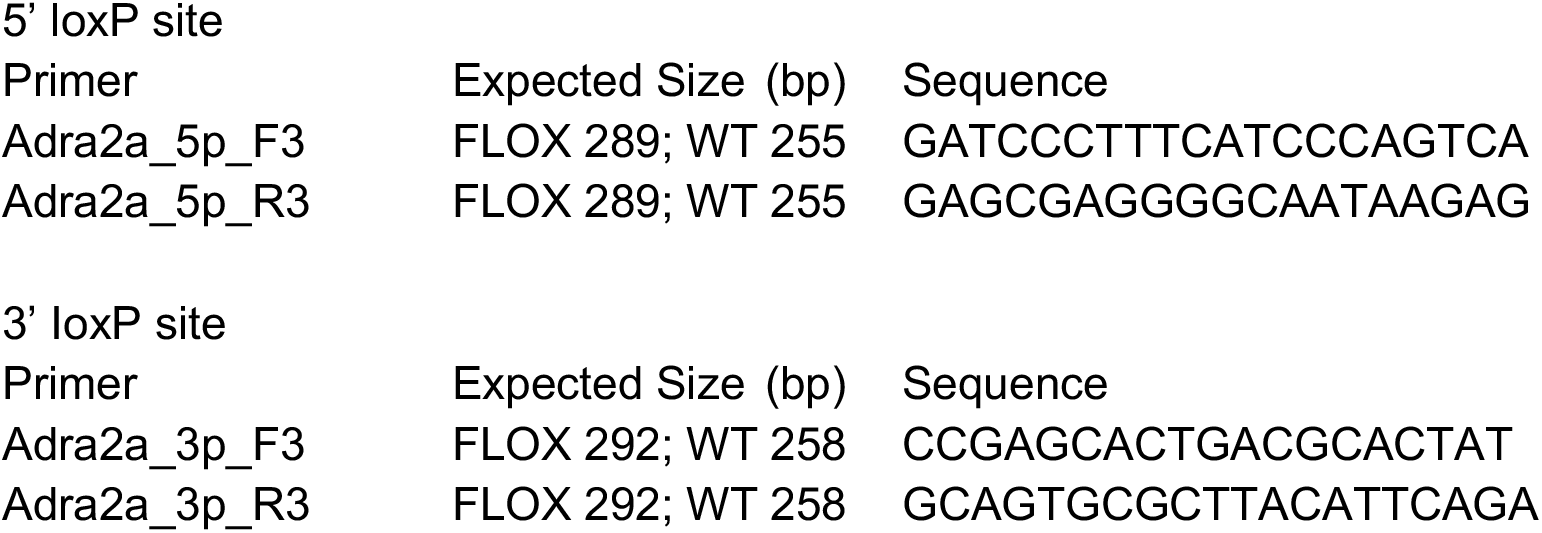

The screen for gene deletion consisted of PCR using the Adra2a_5p_F3 and Adra2a_3p_R3 and evaluating for bands 100-300 bp in size.

Subsequent genotyping of the backcrossed line and subsequent crosses were performed using PCR with the following primers identifying both the presence of 3’ and 5’ LoxP inserts as well as potential gene deletion events.

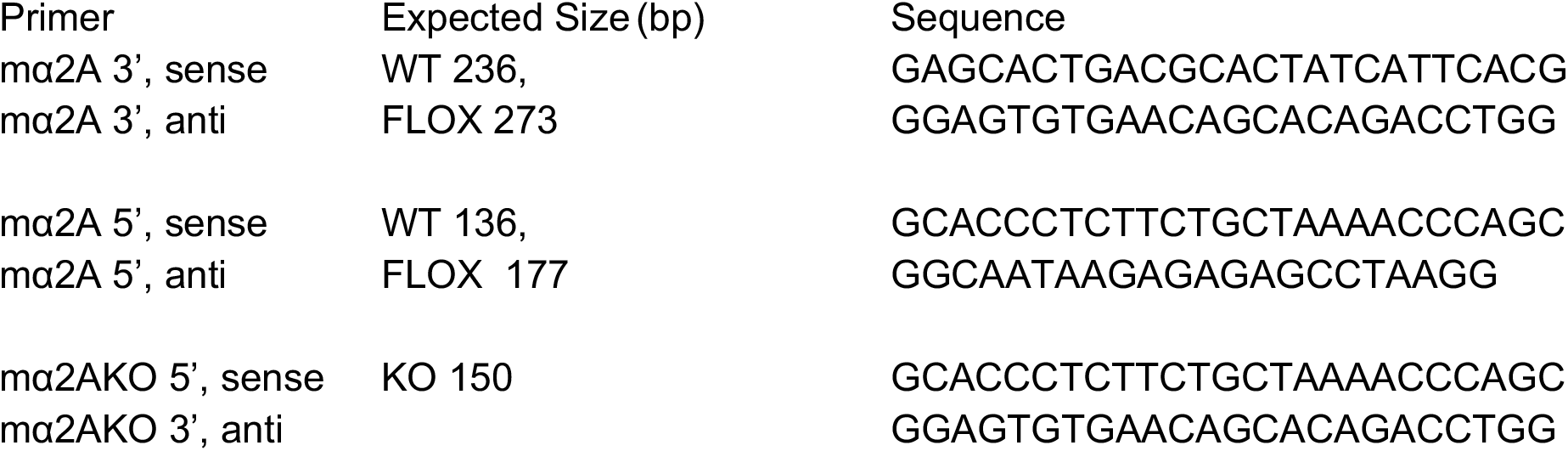

### Fluorescent In-Situ Hybridization

Fresh-frozen brains were imbedded in Tissue-Tek OCT Compound (Electron Microscopy Services, Hatfield PA) cut at 14 μm using a cryostat (Microm HM 505 E Cryostat, Zeiss, Oberkochen Germany) and collected on SuperFrost Plus Slides (Thermo Fisher, Waltham MA). All sections were stored at −80 °C. RNAscope Multiplex Fluorescent Reagent Kit v2 (Advanced Cell Diagnostic, Newark CA) was used for in situ hybridization with the following probes: Mm-Adra2a (Advanced Cell Diagnostic cat. 425341) Mm-Slc17a6-E2-C2 (Advanced Cell Diagnostic cat. 428871-C2), and Mm-Dbh-O1-C3 (Advanced Cell Diagnostic cat. 464612-C3), or Mm-Slc32a1-C3 (Advanced Cell Diagnostic cat. 319191-C3); and the following dyes: TSA Vivid fluorophore 520 (Advanced Cell Diagnostic cat. 323271), TSA Vivid fluorophore 570 (Advanced Cell Diagnostic cat. 323272), and TSA Vivid fluorophore 650 (Advanced Cell Diagnostic cat. 323273). In situ RNA hybridization was performed according to the manufacturer’s instructions, with the following modifications: tissue was fixed in cold 4% paraformaldehyde (PFA) for 60 min prior to serial dehydration and stored overnight at 4 °C in 100% ethanol before proceeding to hybridization. TSA vivid fluorophores were diluted 1:800 prior to application. Images were acquired using a Keyence BZ-X800 (Keyence, Itaska IL). Image processing and analysis was conducted using the Fiji distribution of ImageJ2 and QuPath 0.6.0.^22,23^

### Righting Reflex Assay

Mice were placed in partially-open 200 mL cylindrical chambers on an external heating pad set at 38 °C. Mice were given 1 mg/kg intraperitoneal (IP) dexmedetomidine (dexmedetomidine HCl, Sigma-Aldrich, in PBS) and righting reflex assessed 20 minutes after injection. A mouse was deemed to have an intact righting reflex if it returned to a prone position after being placed supine by rotating the chamber. If the mouse failed to return to the prone position within 10 seconds of being placed supine, the reflex was classified as absent.

### Spontaneous Movement (Beam Break Assay)

Mouse spontaneous ambulation was recorded using a 1-dimensional cage locomotor infrared beam system and accompanying Accuscan Fusion 4.1 software (Omnitech Electronics Inc, Columbus OH). Actigraphy was recorded for a total of one hour starting 30 minutes after IP injection of vehicle, 0.3, or 1.0 mg/kg dexmedetomidine and total distance travelled recorded.

### Forced Movement (Rotarod Assay)

Prior to any testing, all mice underwent habituation on the rotarod (IITC Life Science, Woodland Hills, CA) over 3 training days. On the first day, they were placed on the rotarod at a constant speed of 5 revolutions per minute (RPM) and had to run for at least 60 seconds. The training trial was repeated twice more with at least 5 minutes of rest between trials. On the second day, the same procedure was followed with the revolving speed of the rotarod increased to 10 RPM, and on the third day the speed was increased to 15 RPM. On a testing day, mice received an IP injection of vehicle, 0.3, or 1.0 mg/kg dexmedetomidine, and after 20 minutes were placed on the rotarod where the RPM increased from 5 to 15 over the course of 60 seconds and time to fall recorded (with a maximum time of 60 seconds.) This trial was repeated twice more for a total of three trials with at least 1 minute of rest between trials.

### EEG headpiece implantation surgery

Mice were anesthetized with 2% isoflurane in 100% oxygen for induction, and maintained on 1.5% isoflurane for the surgery. After confirmation of anesthetic induction with lack of response to toe pinch, mice were placed on a stereotaxic frame (Kopf Model 902, Kopf Instruments, Tujunga CA). Eyes were protected by using eye ointment and core temperature was maintained at 37 ± 1.0°C using a closed-loop heating pad (CWE Inc, Ardmore PA).

EEG/EMG headpieces were constructed and implanted as previously described^24,25^ with the following modifications: headpieces consisted of a single midline-spaning 2 × 6 rows of pins, with 10 total epidural EEG leads arranged at 1.3 mm intervals from 2.3 mm anterior to bregma to 2.9 mm posterior to bregma along with two cervical EMG leads. EEG leads were located 0.65 lateral to midline, with leads in each hemisphere. An anchor screw was implanted at 2 mm posterior to bregma and 2.5 mm left-lateral.

### EEG and EMG Acquisition

EEG and EMG were recorded as described previously^25,26^ using Intan headstages (Intan Technologies) through an acquisition box constructed according to Open-Ephys specifications (Open-Ephys, Atlanta GA).^27^ All signals were recorded using the freely available open source open-ephys GUI acquisition software (v1.0, www.open-ephys.org). Mice were attached to headstages in a heated chamber, and allowed to acclimate for 15 minutes before 30 minutes of baseline recording. Mice were then given 0.3 mg/kg IP dexmedetomidine and recording continued for 90 minutes.

### EEG Preprocessing and Analysis

All EEG was recorded using an acquisition system constructed according to open-source designs available from Open-Ephys, 32-channel headstage amplifiers (RHD 2132, Intan Technologies, Los Angeles, CA), and the Open-Ephys GUI software v2.01. Recordings were imported into Matlab 2024b (Mathworks, Natick MA) for post-processing. EEG was bandpass filtered between 1 and 55 Hz using a 6th order zero-phase Butterworth filter prior to additional processing. Impedance measurements were taken prior to recordings and leads with an impedance measurement greater than 30 kΩ were excluded from analysis. All remaining leads were visually inspected for artifacts, and the best quality artifact-free frontal lead from each mouse was used for subsequent analysis. Recordings were further manually examined for artifact and periods with significant artifact excluded from analysis. Analyses were performed on the 30 minutes of baseline recording and the period 20-60 minutes after dexmedetomidine administration (capturing the period of maximum drug effect.) Spectral power estimation was computed as described previously,^26^ using previously published code.^28^ 95% bootstrap CIs of the spectra were computed by independently resampling subjects within each group (1,000 iterations).

### Statistical Analysis

Data were analyzed using Matlab 2024b (Mathworks Inc.) and Prism 10.0 (GraphPad). Statistical significance was assessed at α = 0.05, and parametric testing was employed throughout. A p-value < 0.05 was considered statistically significant for all comparisons. Indications of significance are as follows: * p <0.05; ** p < 0.01; *** p < 0.001; **** p < 0.0001.

## Results

### Generation of Conditional *Adra2a* Mouse Line

Employing CRISPR-Cas9 targeted recombination in single-cell mouse embryos, we aimed to create a conditional *Adra2a* allele. Our design was simplified by *Adra2a* being composed of a single exon 3.7 kB long. Flanking loxP sites were inserted 5’ and 3’ of the untranslated regions (UTR) of the gene to permit efficient excision of the entire coding region by Cre recombinase. In conjunction with the Transgenic and Chimeric Mouse Core of the University of Pennsylvania, we designed sgRNA (to cut at insertion sites when in complex with Cas9) and 160 bp single-stranded oligonucleotide repair templates containing loxP elements for insertion by homology-directed repair (Table 1, Figure 1.) Of 38 mice generated and PCR screened for both loxP insertion and deletion, 14 (36.8%) had at least 1 insertion event, and 12 (31.6%) had a complete or partial gene deletion event, with 3 of the mice with insertions having homozygous insertions at both the 5’ and 3’ sites without deletions. Those 3 mice underwent Sanger sequencing confirming the targeted loxP insertions with a complete, intact coding region of the gene. Using 1 of these 3 founders, the line, *Adra2a*^fl/+^ was then backcrossed into a mixed C57BL/6 and 129/Sv background for 5 generations to eliminate any unrecognized, off-target recombination events. *Adra2a*^fl/+^ mice were then interbred to generate homozygous *Adra2a*^fl/fl^ mice that exhibited normal health and fertility.

**Table 1.**
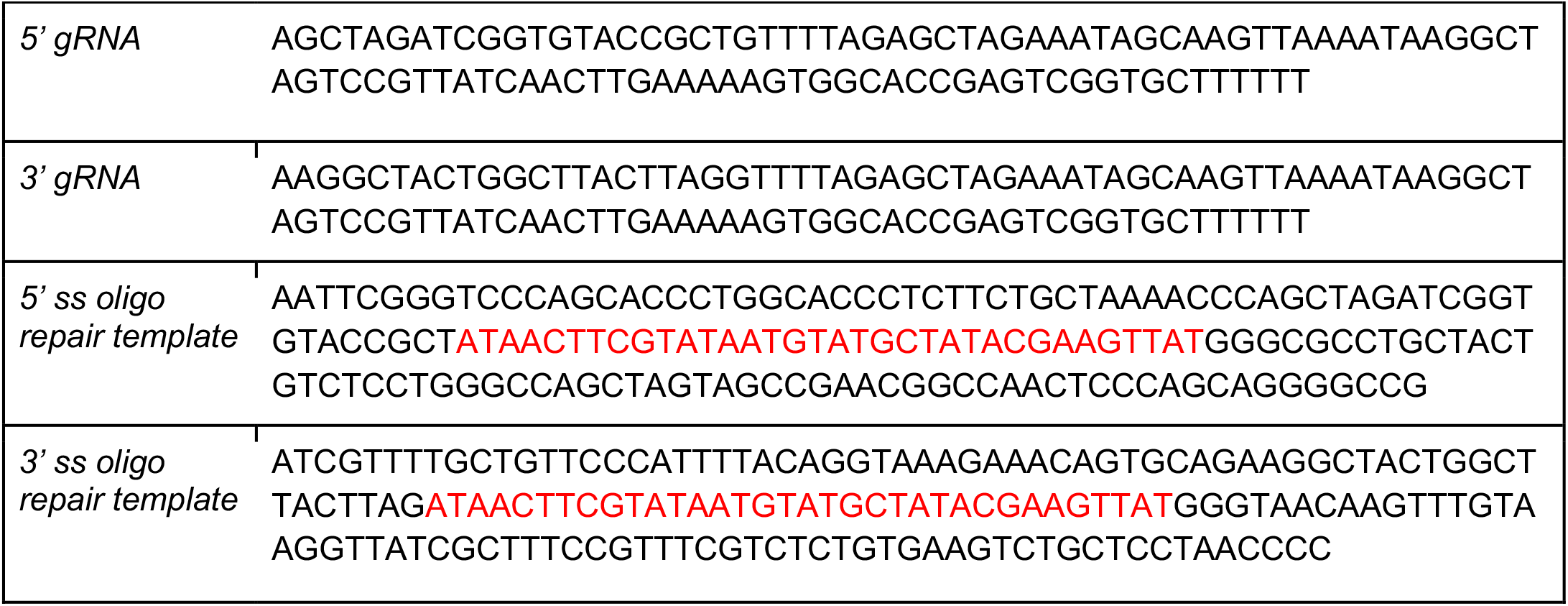
Guide RNA and Single Strand Oligonucleotide Repair Template Sequences (loxP in Red)

**Figure 1.**
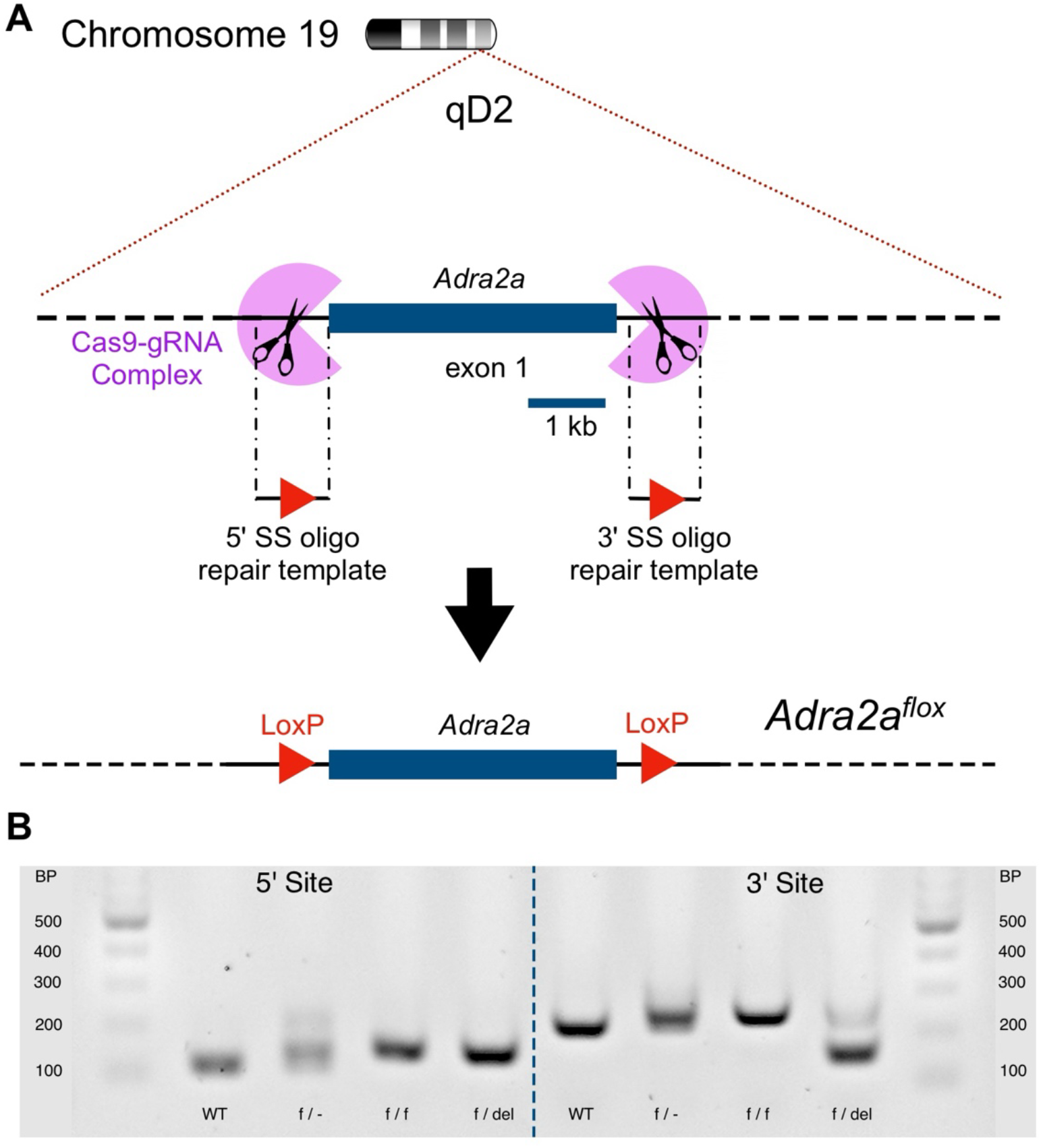
Creation and genotyping of *Adra2a*^flox^, a conditional *Adra2a* mouse line. **(A)** Flanking loxP sites (flox) were introduced around the single exon of *Adra2a* using CRISPR/Cas9. **(B)** Example PCR genotyping gel for the *Adra2a*^flox^ line. At the 5’ site, the expected WT band is 136 bp, while the floxed band is 177 bp. For the 3’ site, the WT band is 236 bp while the floxed band is 273 bp. Gene deletion generates a 150 bp band

### Genomically-expressed Cell-type-specific *Cre* Expression in *Adra2a^fl/fl^* Mice Produces Cell-type-specific Adra2a-Knockout

We crossed our *Adra2a*^fl/fl^ line with mouse lines possessing genomically-encoded Cre recombinase under cell-specific promoters, including GABAergic (*Slc32a1* for VGAT), adrenergic (*Dbh*), and pan-neuronal (*Snap25*) promoters. To confirm cell-specific deletion of *Adra2a*, the presence of its transcript, as well as those of cell-specific markers, were evaluated in the brains of *Adra2a*^fl/fl^ mice and Cre-expressing crosses using fluorescent in-situ hybridization (Figure 2A-H). Analysis of *Adra2a* expression in the region of the locus coeruleus and the preoptic area, areas with known wild-type expression of *Adra2a*, showed profound transcript reduction in cell types with expected Cre expression (Figure 2I, J).

**Figure 2.**
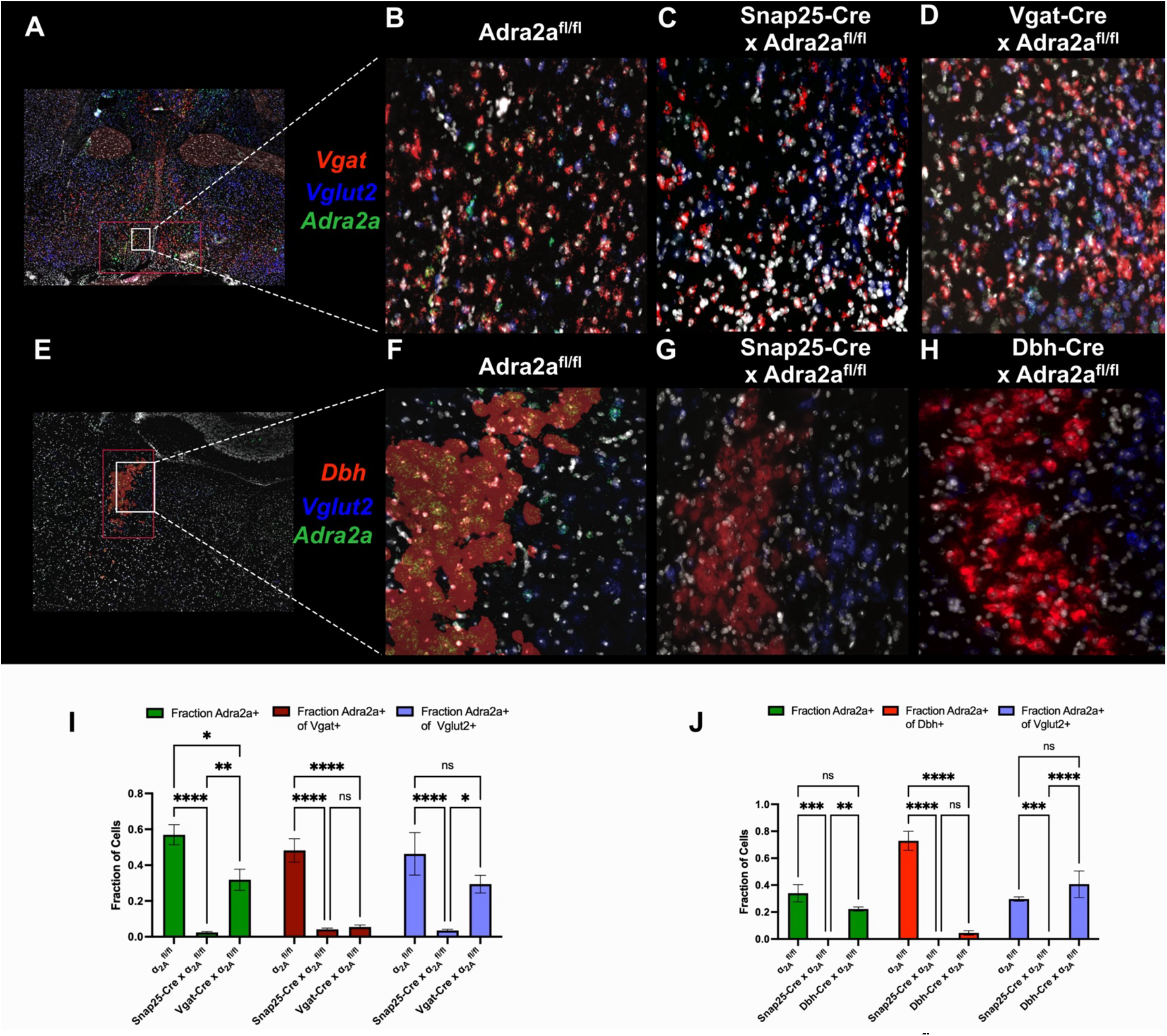
Fluorescent *in situ* hybridization shows cell-specific knockout in *Adra2a*^flox^ crosses in lines with Cre under cell-specific promoters. FISH in the preoptic area (POA (**A-D**); bregma +0.02 mm) and locus coeruleus (LC (**E-H**) bregma −5.40 mm) demonstrates cell-type-selective knockout of *Adra2a* in crosses with Cre driven by *Snap25* (**B**,**G**) *Slc32a1* (VGAT) (**D**), and *Dbh* (**H**). (**I**) In the POA, the *Snap25*-Cre cross shows a significant reduction in the fraction of cells expressing *Adra2a*, while the *Vgat*-Cre line only shows a significantly reduced fraction of *Adra2a* of *Slc32a1*-expressing cells, but not in *Slc17a6* (VGluT2)-expressing cells. (**J**) Similarly, in the region of the LC, the *Snap25*-Cre cross shows a significant reduction in fraction of cells expressing *Adra2a*, and the *Dbh*-Cre cross line only shows a significantly reduced fraction of *Adra2a* of *Dbh*-expressing cells, but not in *Slc17a6*-expressing cells. (n=3 per genotype, 2-way ANOVA, *p<0.05,**p<0.01, ***p,0.001, ****p<0.0001)

### Neuronal Knockout of Adra2a Confers Broad Resistance to Central Alpha-2 Agonist Effects, while Adrenergic and GABAergic Knockout Mice Show Resistance to Distinct Aspects of Sedation

Neither the pan-neuronal *Adra2a* knockout nor the adrenergic *Adra2a* knockout lost the righting reflex– a rodent behavioral proxy for loss of consciousness– at a hypnotic dose of the α_2_ agonist dexmedetomidine, while the GABAergic *Adra2a* knockout showed similar sensitivity to the floxed control (Figure 3.) While at baseline, all lines had similar basal temperatures (Figure 4A) the pan-neuronal *Adra2a* knockout showed complete resistance to another centrally-mediated α_2_ effect, hypothermia, while the adrenergic-specific knockout showed some resistance to this effect at higher doses and the GABAergic knockout was again indistinguishable from floxed controls (Figure 4B). We then examined the effect of cell-specific knockouts of the *Adra2a* gene on two behavioral measures of sedation, spontaneous locomotion (Figure 5) and coordination with forced movement (Figure 6). While all mice behaved similarly with saline control injections, the pan-neuronal *Adra2a* knockout showed complete resistance to central effects of dexmedetomidine in both behavioral measures of sedation. Adrenergic and GABAergic *Adra2a* knockouts diverged in their resistances, with GABAergic knockouts resisting only the decrease in spontaneous movement elicited by dexmedetomidine compared to floxed controls (Figure 5), and adrenergic knockouts only resisting impaired coordination with forced movement (Figure 6).

**Figure 3.**
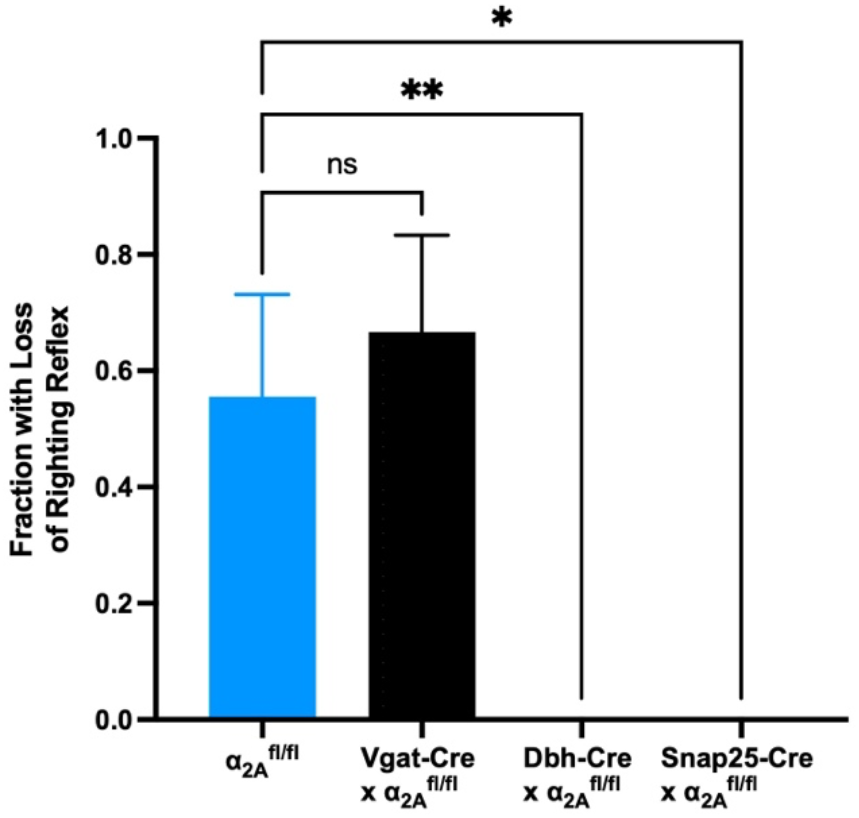
Pan-neuronal and adrenergic *Adra2a* knockout are resistant to dexmedetomidine-induced hypnosis as measured by loss-of-righting-reflex (LoRR). Mice were given 1 mg/kg dexmedetomidine IP and righting reflex assessed after 20 minutes. *Snap25*-Cre and *Dbh*-Cre *Adra2a*^*flox*^crosses showed complete resistance to LoRR, while the sensitivity of the *Vgat*-Cre cross did not vary significantly from floxed controls. (One-way ANOVA, n=9-18 per group. *p<0.05, **p<0.01).

**Figure 4.**
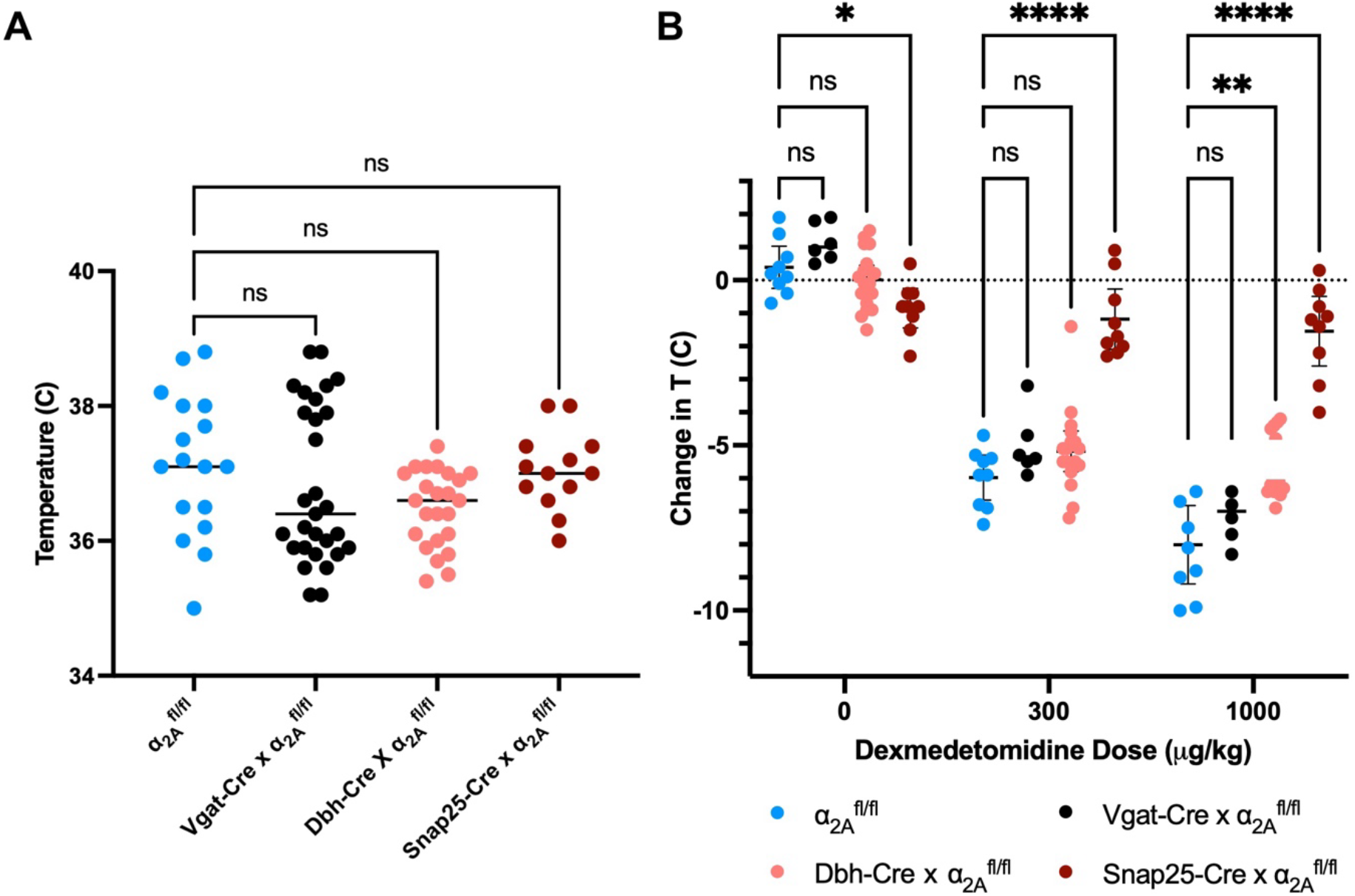
Baseline temperatures and temperature change with α_2_-agonist administration by genotype. **(A)** Core temperatures did not vary significantly by genotype. (One-way ANOVA, n=13-29 per group) **(B)** Core temperature was assessed 20 minutes after IP dexmedetomidine at 0, 300, or 1000 μg/kg; data are expressed as delta temperature (post − pre administration). *Snap25*-Cre x *Adra2a*^fl/fl^ mice showed complete resistance to α_2_-agonist-induced temperature change, while the *Dbh*-Cre cross showed some resistance at the highest dose. (2-way ANOVA, n=6-18 per group. *p<0.05, **p<0.01, ****p<0.0001)

**Figure 5.**
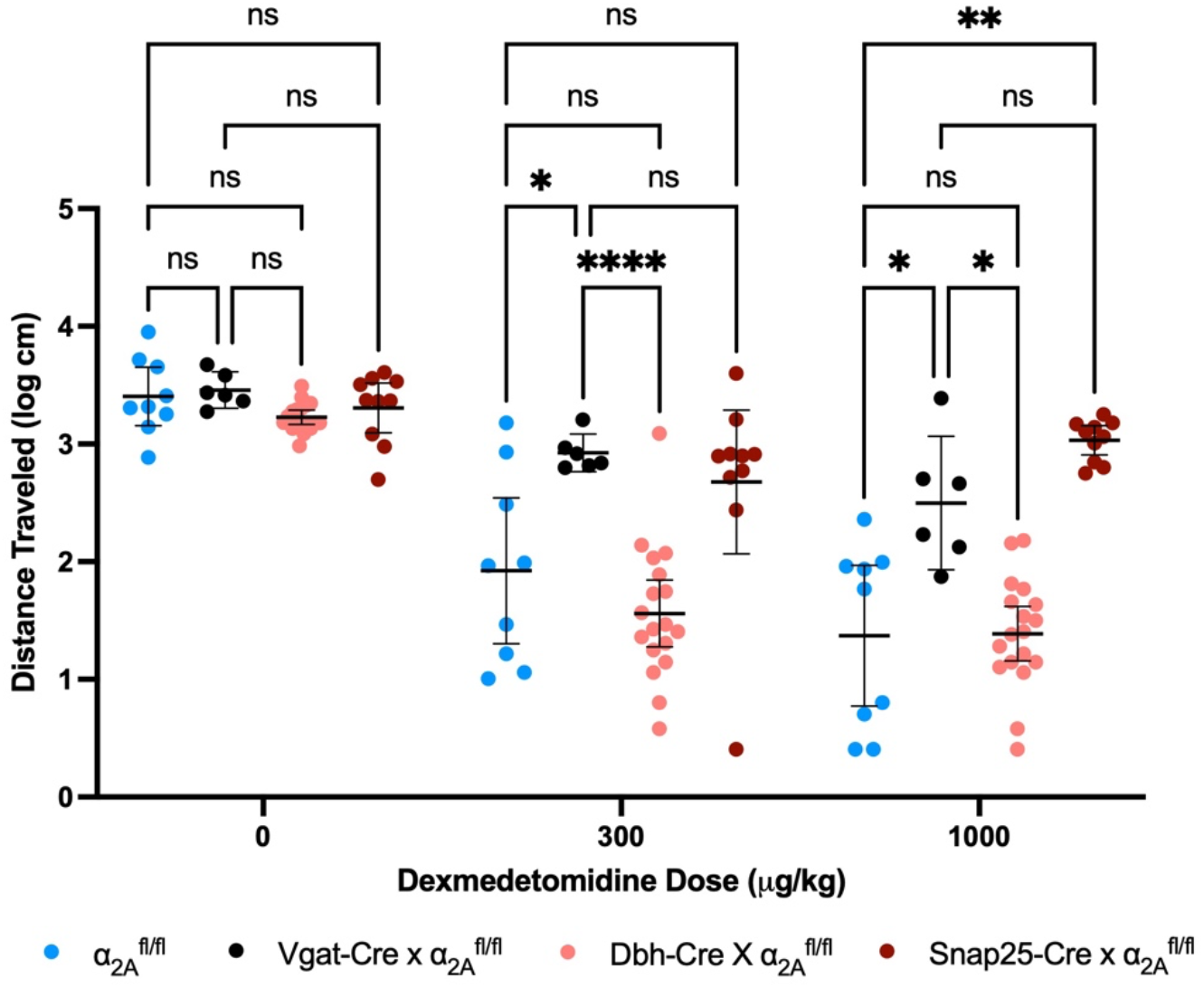
Spontaneous movement inhibition with α_2_-agonist administration. Spontaneous movement was measured by beam break 30-90 minutes after drug administration. The pan-neuronal *Adra2a* knockout showed significant resistance to α_2_-agonist reduction in distance traveled, as did the GABAergic-specific knockout. *Dbh*-specific *Adra2a* knockouts were indistinguishable from floxed controls. (2-way ANOVA. n=6-18 per group. *p<0.05, **p<0.01, ****p<0.0001)

**Figure 6.**
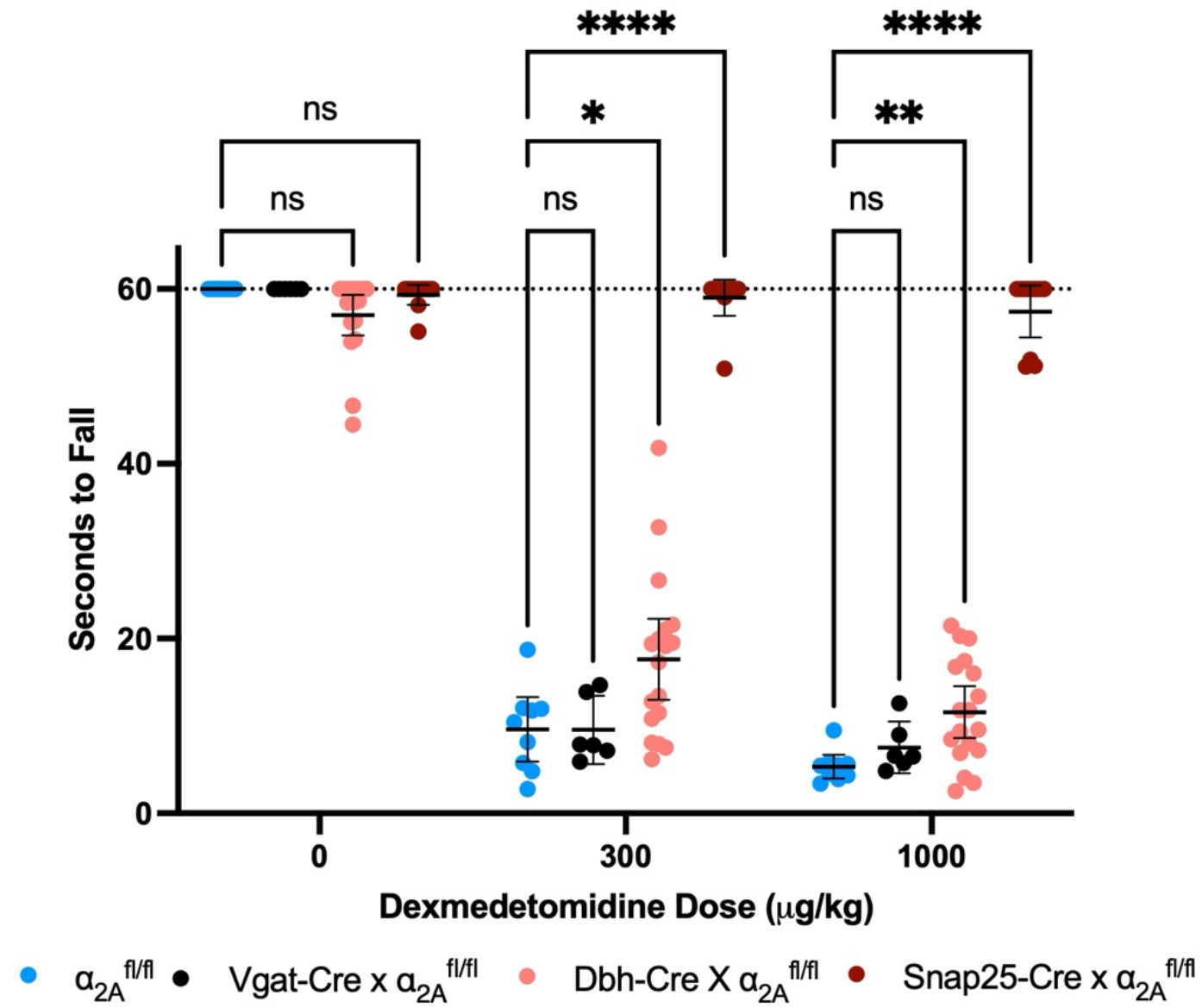
α_2_-agonist effects on coordination with forced movement by genotype. Coordination with forced movement was assessed via rotarod 20 minutes after dexmedetomidine injection. *Snap25*-Cre x *Adra2a*^fl/fl^ mice showed no inhibition of coordination with dexmedetomidine, *Dbh*-Cre x *Adra2a*^fl/fl^ mice showed some resistance, and *Vgat*-Cre crosses were indistinguishable from floxed controls. (2-way ANOVA. n=6-18 per group. *p<0.05, **p<0.01, ****p<0.0001)

### EEG Spectral Changes in Response to Alpha-2 Agonist Reflect Behavioral Drug Effects

Among baseline EEG across the lines, only the GABAergic *Adra2a* knockout line showed a baseline difference in normalized spectra from controls, with a higher proportion of delta power and less low beta power, despite similar behavior at baseline (Figure 7A,C). At a sedative dose of dexmedetomidine (0.3 mg/kg IP) floxed controls as well as adrenergic and GABAergic *Adra2a* knockouts showed increases in the proportion of delta power relative to baseline (Figure 7B,D). Adrenergic knockouts further showed a decrease in relative alpha power and an increase in a proportion of beta power (∼18-25 Hz). In contrast, the pan-neuronal *Adra2a* knockout showed no change in its EEG spectrum with dexmedetomidine administration (Figure 7B,D).

**Figure 7.**
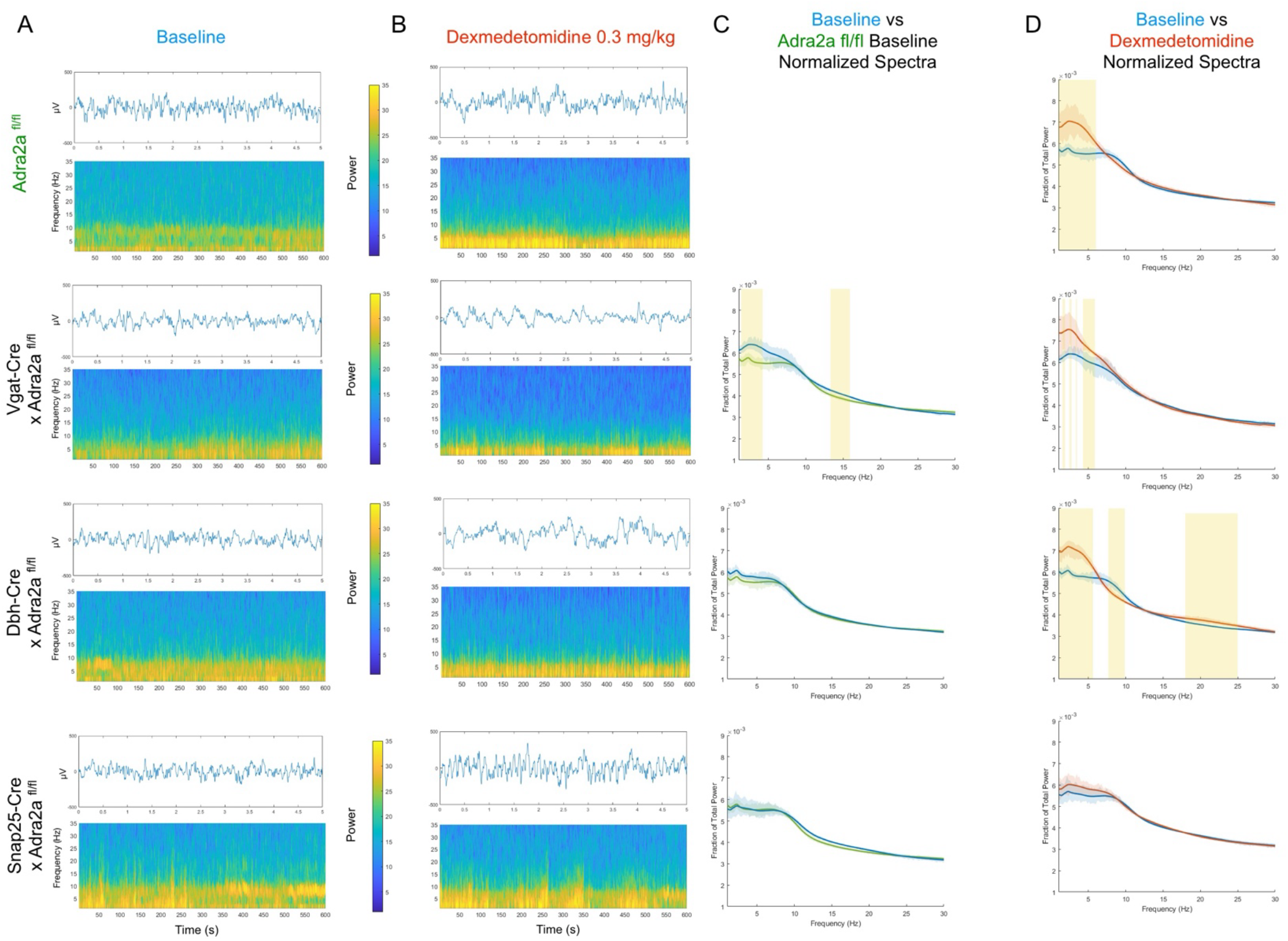
Pan-neuronal *Adra2a* knockouts do not show changes to EEG with dexmedetomidine administration. (**A**) EEG was recorded for 30 minutes at baseline. Example raw (top, 5 seconds shown) and spectrogram (bottom, 10 minutes shown) plots are shown for each genotype. (**B**) Similar examples of the same genotypes at 20-60 minutes after dexmedetomidine administration (0.3 mg/kg, i.p.). (**C**) At baseline, the spectra of various cell-specific *Adra2a* knockouts were largely indistinguishable from floxed control. *Vgat*-Cre x *Adra2a*^fl/fl^ mice however showed an elevated proportion of delta power and reduced low beta power relative to floxed controls (mean spectra and bootstrapped 95% CI shown, n=4-5 per genotype, yellow highlights indicate areas where 95% CI do not overlap.) (**D**) Floxed controls, *Dbh*-Cre crosses, and *Vgat*-Cre crosses all show increases in their proportion of delta power following dexmedetomidine administration, with the Dbh-Cre cross also showing a relative decrease in alpha power and increase in beta power. The Snap25-Cre cross, in contrast, shows no spectral changes with dexmedetomidine administration (mean spectra and bootstrapped 95% CI shown, n=4-5 per genotype, yellow highlights indicate areas where 95% CI do not overlap).

## Discussion

We have created a conditional *Adra2a* mouse line through the insertion of flanking loxP sites 5’ and 3’ of the UTRs for the gene, allowing for genetic dissection of the role of adrenergic α_2A_-receptors. We confirmed specificity of conditional knockouts through crosses of our floxed line with other established lines genomically expressing Cre-recombinase under cell-specific promoters and using *in situ* hybridization to demonstrate selective knockdown of *Adra2a* mRNA in the relevant cell types. The pan-neuronal *Adra2a* knockout showed profound resistance to known centrally-mediated effects of α_2_-agonists, including hypothermia, sedation, and hypnosis. In parallel to that finding of resistance, the pan-neuronal knockout did not show significant changes to its EEG spectrum with administration of the α_2_-agonist dexmedetomidine. Together, these show that α_2A_-receptors on neurons are necessary for centrally-mediated effects of α_2_-agonists. While our findings do not rule out a role for non-neuronal (astrocytic and microglial) cells in central α_2A_ effects, they do show that α_2A_ receptors on astrocytes and glia are not sufficient to drive those effects. Adrenergic- and GABAergic-specific *Adra2a* knockouts demonstrate resistance to mutually exclusive measures of behavioral sedation in response to the α_2_-agonist dexmedetomidine. GABAergic-specific knockouts do not exhibit as great a decrease in spontaneous movement as floxed controls when given dexmedetomidine, while adrenergic-specific knockouts show resistance to impairment of coordination with forced movement with α_2_-agonist administration. In addition, adrenergic-specific knockouts demonstrate some resistance to the hypothermic effects of dexmedetomidine. EEG spectra for both adrenergic- and GABAergic-knockouts meanwhile show increases in the proportion of delta power with α_2_-agonist administration, consistent with classic signatures of sedation on EEG.^29^ Together, this suggests that separate specific behaviors induced by α_2_-agonists that are typically grouped under the umbrella of sedation, reduction of spontaneous movement and impaired coordination with forced movement, are mediated through α_2A_-receptors on GABAergic and adrenergic neurons, respectively.

The divergence in behavioral phenotype between the GABAergic- and adrenergic-specific *Adra2a* knockouts with dexmedetomidine administration is unexpected and demonstrates a pitfall of using overbroad descriptors like “sedation.” In the adrenergic-specific knockouts, volitional locomotor activity is unchanged compared to controls, while coordination and hypnosis (as assessed by loss-of-righting-reflex) effects in response to dexmedetomidine are abrogated. This could be consistent with a non-specific functional lessening of sedative-hypnotic potency–as volitional effects are most sensitive to sedative-hypnotics, with coordination following, and finally consciousness requiring the highest dose of sedative-hypnotics to ablate.^30^ The relative preservation of volitional movement even with loss of coordinated movement seen in the GABAergic *Adra2a* knockout, in contrast, does not conform to the standard order of loss of function with sedative-hypnotics. Impaired function in prefrontal α_2A_ receptors is known to be associated with locomotor hyperactivity,^31^ though we do not see such increased activity at baseline in the GABAergic knockouts. Given the presence of α_2A_ receptors on both prefrontal pyramidal neurons as well as prefrontal GABAergic interneurons, it is possible that an imbalance in the excitation/inhibition with the addition of an α_2_ agonist could trigger hyperlocomotion, as decreased prefrontal glutamatergic activity has also been implicated in a hyperactivity phenotype.^32^ Future virally-based locational and genetic-specific knockouts using the conditional *Adra2a* knockout line will be better equipped to answer this specific question.

Despite a greater than 90% reduction across the board in the proportion of cells with *Adra2a* mRNA that putatively produce Cre recombinase, we did not see a complete elimination with our *in situ* hybridization studies. This is likely due to the limitations of the imaging resolution employed rather than aberrant expression of Cre or failure of *Adra2a* knockout in the presence of Cre. In the *Snap25*-Cre crosses, microglia and astrocytes would be expected to continue to express *Adra2a*. Even with our high-resolution and magnification fluorescence microscopy, it is occasionally difficult to distinguish closely apposed cells even with sections cut at roughly the width of a cell, 14 μm. This could lead to glial expression of Adra2a visually overlapping with neuronal expression of *Slc32a1, Slc17a6*, or *Dbh*. While there is evidence that some of these other markers may rarely be expressed in non-neurons,^33^ visual overlap seems the more likely culprit. Relatedly, in the *Slc32a1*-Cre crosses, in the rare cases where we see *Slc32a1* and *Adra2a* expression, it frequently also coincides with *Slc17a6* expression. While there are neurons that produce both GABA and glutamate,^34^ the simpler and thus more likely explanation of such apparent exclusive triple-expression in putative cell-selective knockouts is that a *Slc17a6-* and *Adra2a*-expressing neuron is sitting directly next to or on top of a *Slc32a1*-expressing neuron.

The lines that demonstrated some degree of behavioral sedation in response to α_2_ agonist administration also showed an increase in the proportion of EEG delta power in response to drug administration, as would be expected with α_2_ agonist sedation. Though the floxed control and cell-specific *Adra2a* knockout lines were behaviorally indistinguishable from each other at baseline, the GABAergic-specific knockout did demonstrate a higher proportion of delta power at baseline compared to the floxed control. This difference at baseline could be due to differential cortical activity primarily in the prefrontal cortex, where α_2A_ receptors have been shown to be present in layer 6 (putatively GABAergic),^35^ and on confirmed GABAergic interneurons.^36^ Alternatively, it could arise from differential architecture from altered GABAergic neuron migration during development, to which *Adra2a* contributes.^37^ To distinguish whether this altered baseline activity is an artifact of development, future studies could use an inducible *Vgat*-driven Cre^38^ to selectively knockout *Adra2a* in *Slc32a1*-expressing neurons in adult animals, avoiding developmental effects.

Characteristics of our adrenergic-specific *Adra2a* knockout line, in terms of hypnotic and temperature effects, are consistent with previous studies of α_2A_ receptors. Early studies of the central effects of α_2_ agonists also found that righting reflex could be restored with local application of an α_2_ antagonist at the locus coeruleus, the site of 75% of central adrenergic neurons.^6^ RNAi-mediated knockdown of *Adra2a* in the locus coeruleus was similarly able to confer resistance to loss-of-righting reflex, while not affecting behavioral measures of sedation, or EEG-related changes.^8^ Relatedly, the locus coeruleus plays a key role in sensory-mediated awakenings from sleep, with increased locus activity decreasing the sensory stimulus needed to wake and decreased activity raising the threshold.^39^ Righting reflex testing may serve as a rough vestibular evoked response, and preserving the activity of the locus coeruleus through eliminating the effect of α_2_ agonist effect on those cells could preserve that response. Finally, a recent report demonstrated the role of adrenergic neurons in maintaining temperature during dexmedetomidine sedation,^40^ which is also congruent with our finding that adrenergic-specific *Adra2a* knockouts maintain their temperatures compared to either floxed controls or the GABAergic-specific knockouts.

Our finding of differential cell-specific α_2A_ effects on volitional locomotion and coordinated movement demonstrates the potentially complex and dissociable nature of adrenergic signaling on multiple arms of a central process. The behavioral divergence between GABAergic- and adrenergic-specific *Adra2a* knockouts suggests that that α_2_-agonist sedation is a multi-site process with distinct receptor populations governing specific behavioral components of the phenomena. The creation of the conditional *Adra2a* knockout opens the door to a host of investigations into adrenergic signaling and function. Beyond sedation, the conditional line provides a platform for studying the contributions of cell-type-specific adrenergic signaling to arousal, cognition, pain modulation, stress responses, and cardiovascular regulation. Ultimately, this work lays a genetic and conceptual foundation for dissecting the full complexity of α_2a_-adrenergic neurobiology.

## Acknowledgements

We are grateful to the Transgenic and Chimeric Mouse Core of the University of Pennsylvania for their assistance in generation of the *Adra2a*^fl/fl^ line (supported by NIH center grants P30DK050306, P30DK019525, and P30CA016520).

